# PTEN deletion in spinal pathways via retrograde transduction with AAV-rg enhances forelimb motor recovery after cervical spinal cord injury; sex differences and late-onset pathophysiologies

**DOI:** 10.1101/2023.03.20.533502

**Authors:** Mariajose Metcalfe, Oswald Steward

## Abstract

Spinal cord injuries (SCI) cause permanent functional impairments due to interruption of motor and sensory pathways. Regeneration of axons does not occur due to lack of intrinsic growth capacity of adult neurons and extrinsic inhibitory factors, especially at the injury site. However, some regeneration can be achieved via deletion of the phosphatase and tensin homolog (PTEN) in cells of origin of spinal pathways. Here, we deployed an AAV variant that is retrogradely transported (AAV-rg) to deliver gene modifying cargos to the cells of origin of multiple pathways interrupted by SCI, testing whether this promoted recovery of motor function. PTEN^f/f^;Rosa^tdTomato^ mice and control Rosa^tdTomato^ mice received injections of different doses (number of genome copies, GCs) of AAV-rg/Cre into the cervical spinal cord at the time of a C5 dorsal hemisection injury. Forelimb grip strength was tested over time using a grip strength meter. PTEN^f/f^;Rosa^tdTomato^ mice with AAV-rg/Cre (PTEN-deleted) exhibited substantial improvements in forelimb gripping ability in comparison to controls. Of note, there were major sex differences in the extent of recovery, with male mice exhibiting greater recovery than females. However, at around 5-7 weeks post-injury/injection, many mice with SCI and AAV-rg-mediated PTEN deletion began to exhibit pathophysiologies involving excessive scratching of the ears and back of the neck and rigid forward extension of the hindlimbs. These pathophysiologies increased in incidence and severity over time. Our results reveal that although intra-spinal injections of AAV-rg/Cre in PTEN^f/f^;Rosa^tdTomato^ mice can enhance forelimb motor recovery after SCI, late-developing functional abnormalities occur with the experimental conditions used here. Mechanisms underlying late-developing pathophysiologies remain to be defined.

## 1. INTRODUCTION

Spinal cord injuries (SCI) interrupt descending motor and autonomic pathways and ascending sensory pathways in the spinal cord, causing paralysis, loss of sensation and loss of bladder and bowel function. Damaged connections within the spinal cord do not regenerate and the prevailing concept is that regeneration failure is due to lack of intrinsic growth capacity of adult neurons and extrinsic inhibitory factors present in the mature nervous system, especially at the injury site.

Neuron-intrinsic signaling pathways that enable a regenerative response have been identified. In this context, the phosphoinositide 3-kinase (PI3K)/mechanistic target of rapamycin (mTOR) pathway plays an important role for regulating cell size, protein synthesis, survival and cytoskeleton formation necessary for axon extension after injury (Leibinger et al., 2019; Proud, 2002; Testa and Tsichlis, 2005; Zhou and Snider, 2006). Activation of PI3K via receptor tyrosine kinase (RTK) leads to activation of intracellular serine/threonine protein kinase AKT and downstream activation of the mTOR pathway. Activation of mTOR leads to the phosphorylation of different substrates, including ribosomal protein S6 (pS6), one consequence of which is increased translation of 5’ TOP mRNAs important for cell growth (Guertin and Sabatini, 2007). There are other important consequences of mTOR activation in general and S6 phosphorylation in particular including alterations in gene transcription in the nucleus (Proud, 2002).

An important negative regulator of the AKT/mTOR pathway is phosphatase and tensin homolog (PTEN), a phosphatase that converts inositol triphosphate (InsP3) to inositol biphosphate (IP2) reversing the action of PI3K. Overall activation of mTOR depends on the balance between PI3K and PTEN, so PTEN inactivation drives the mTOR pathway.

Studies from multiple labs have used adeno-associated virus (AAVs) to drive mTOR activity by genetic deletion or knockdown of PTEN delete to enable axon regeneration. AAVs are an effective platform for *in vivo* gene therapy, as they mediate high levels of transgene expression in non-dividing cells, are non-toxic and present low innate immunity (Kaplitt et al., 2007). The initial landmark study deployed AAV2/Cre in PTEN^f/f^ mice to delete PTEN in retinal ganglion cells, documenting regeneration with optic nerve crush (Park et al., 2008). Then using PTEN^f/f^ mice and injecting AAV2/Cre into the sensorimotor cortex, un-precedented regeneration of CST axons was documented after SCI (Liu et al., 2010). This has been confirmed in multiple follow-up studies (Danilov and Steward, 2015; Geoffroy et al., 2016; Liu et al., 2010; Willenberg et al., 2016). Of note, the regeneration-enabling effect of PTEN deletion is blocked by rapamycin, implicating mTOR as the critical molecular signaling pathway (Park et al., 2010). Knockdown of PTEN via short-hairpin RNA (shRNA) in the sensorimotor cortex, also enables regeneration of CST axons and enhances motor recovery after injury (Lewandowski and Steward, 2014; Zukor et al., 2013).

These studies document that AAV-based delivery of cargos can be effective in enabling regeneration after SCI, but the strategy has been to target one pathway at a time. For example, regeneration of CST axons is enabled by targeting the sensorimotor cortex (Liu et al., 2010) and regeneration of rubrospinal tract axons by targeting the red nucleus (Challagundla et al., 2015). This one pathway at a time approach does not address the critical unmet need for regeneration of multiple pathways following SCI.

A potential tool to deliver gene-modifying cargos to the cells of origin of multiple spinal pathways in widely-distributed brain regions comes from the development of the retrogradely transported AAV variant: AAV-rg (Tervo et al., 2016). When injected into the spinal cord, AAV-rg is retrogradely transported to the cells of origin of multiple spinal pathways (Tervo et al., 2016) (Metcalfe et al., 2022; Wang et al., 2018). Here we carried out an initial test of whether AAV-rg-based deletion of PTEN in cells of origin of multiple pathways promotes recovery of motor function. For this, we injected AAV-rg/Cre into the cervical spinal cord in PTEN^f/f^;Rosa^tdTomato^ mice and control Rosa^tdTomato^ mice at the time of a C5 dorsal hemisection injury. Forelimb gripping ability was assessed with a grip strength meter. PTEN^f/f^;Rosa^tdTomato^ mice exhibited greater improvements in forelimb gripping ability in comparison to controls. However, at around 5-7 weeks post-injury, some mice began to exhibit pathophysiologies involving excessive scratching and hindlimb dystonia with persistent extension, which increased over time. To begin to assess causes of pathophysiologies, additional studies were carried out to define conditions leading to pathophysiologies. Pathophysiologies were not observed in PTEN^f/f^;Rosa^tdTomato^ mice that received AAV-rg/Cre without concurrent SCI or in PTEN^f/f^;Rosa^tdTomato^ mice that received injections of AAV2/Cre in conjunction with SCI (not AAV-rg/Cre). These results indicate that the pathophysiologies are not due to AAV-rg/Cre mediated deletion alone or selective deletion of PTEN in the spinal cord in conjunction with SCI.

Our findings document that although intra-spinal injections of AAV-rg/Cre in PTEN^f/f^;Rosa^tdTomato^ mice can lead to initial improvements in forelimb motor recovery after SCI, there are late-developing functional abnormalities with the experimental conditions used here. The mechanisms underlying late-developing pathophysiologies remain to be defined.

## 2. MATERIALS AND METHODS

### 2.1 Experimental Animals

#### 2.1.1. Animal use and care

All procedures involving animals were approved by the Institutional Animal Care and use Committee (IACUC) of the University of California Irvine in compliance with the National Institute of Health guidelines. Mice were maintained on a 12h light/dark cycle at 25°C. All procedures were done during the light portion of the cycle.

#### 2.1.2. Mice

Our studies were done using two lines of transgenic mice that we developed in our lab (Gallent and Steward, 2018). One line was created by crossing Rosa26^tdTomato^ mice obtained from the Jackson Laboratory (JAX: 007905) with PTEN^f/f^ mice from our breeding colony to create a new transgenic strain that was homozygous at both loci (PTEN^f/f^;Rosa^tdTomato^). PTEN^f/f^;Rosa^tdTomato^ mice have a lox-P flanked STOP cassette in the ROSA locus and lox-P flanked exon 5 of PTEN so AAV-mediated expression of Cre recombinase deletes PTEN and induces robust expression of tdTomato (tdT) in the same neurons. In creating this new transgenic strain, we also selected for homozygous Rosa^tdTomato^ mice as controls. Because the resulting Rosa^tdTomato^ mice were derived from the crossing, they have a different genetic background from the original, Rosa^tdTomato^ mice obtained from JAX Labs (genetic background B6;129S6).

#### 2.1.3. AAV vectors

To retrogradely transduce neurons in the brain that project to the spinal cord, we used different AAV-rg vectors in different studies. In studies 1&3 (below), AAV-rg/Cre (Addgene viral prep #24593-AAVrg) was used in PTEN^f/f^;Rosa^tdTomato^ mice and control Rosa^tdTomato^ mice. In study 2, which used only PTEN^f/f^;Rosa^tdTomato^ mice, AAV-rg/Cre was used to delete PTEN controls received AAV-rg/GFP (Addgene viral prep #37825-AAVrg). In study 4, mice received AAV-rg/shPTEN/GFP to delete PTEN and controls received similar injections of AAV-rg/Cre. Custom AAV-rg/shPTEN/GFP viral constructs are produced at the University of Pennsylvania Vector Core. The shRNA targeting PTEN has been previously characterized (Lewandowski and Steward, 2014) and it is driven by a human U6 promoter, while, GFP expression is driven by a cytomegalovirus (CMV) promoter. To transduce cells locally within the spinal cord with minimal retrograde transport, Study 5 used AAV2/Cre (Vector Biolabs viral prep #7011).

### 2.2. Spinal cord injuries

Surgical procedures for intra-spinal cord AAV-rg and AAV2 injections at C5 were as previously described (Steward et al., 2021). Mice were anesthetized with Isoflurane (2-3%), their eyes were swabbed with Vaseline, and the skin at the incision site was shaved and cleaned with Betadine. A laminectomy was performed at C5 to expose the underlying spinal cord. Injections (0.3 µl per side) were made using a Hamilton microsyringe fitted with a pulled glass micropipette at 0.5 mm lateral to the midline and 0.5 mm deep.

C5 dorsal hemisections were performed under the same procedure as intraspinal AAV-rg or AAV2 injections. To create a dorsal hemisection, a 15-degree angled Microscalpel (Electron Microscopy Sciences #72046-15) was inserted into the spinal cord at the midline to a depth of 0.5 mm and drawn laterally, then the blade was turned around and drawn in the opposite direction. This was repeated several times to assure completeness of the dorsal transection. After the spinal cord lesion was created, the mouse was passed to a separate station for AAV injections. After completion of the hemisection and intra-spinal cord AAV injections, muscle layers were sutured and the skin was secured with wound clips.

### 2.3. Post-operative care

Following the surgeries, mice were immediately placed on water-circulating jacketed heating pads at 37°C until they fully recovered from the anesthetic. When able to move around in their cage and feed themselves, mice were housed 2-4 per cage on Alpha-Dri bedding. For 3-5 days post-injury, mice received lactated Ringer solution (1ml/20g, subcutaneously) for prophylactic treatment against urinary tract infections. Mice were monitored twice daily for 2 weeks post-injury for general health, coat quality (indicative of normal grooming activity) and mobility within the cage. After this post-operative monitoring, mice were examined by researcher staff at the time of weekly GSM testing and routinely monitored by vivarium staff.

### 2.4. Order of studies to test recovery of function

The overall study of consequences of PTEN deletion on recovery following spinal cord injury (SCI) includes 3 separate studies carried out at different times. Each study involved delivery of different doses of AAV-rg/Cre [dose = number of genome copies (GC) injected per mouse]. For each study, surgeries and AAV-rg/Cre injections were done by the same surgical team and were blind with respect to strain and vector. However, different experimenters were responsible for the functional analyses in each study.

Study 1 was performed to determine if PTEN deletion via intra-spinal injection of AAV-rg/Cre would promote functional recovery after SCI. Study 1 involved 15 PTEN^f/f^;Rosa^tdTomato^ mice (n=11 male; n=4 female) and 14 control Rosa^tdTomato^ mice (n=4 male; n=10 female). Mice were 11 weeks old at the start of the study and were handled for 2 weeks. GSM testing began when mice were 13 weeks of age and continued for 4 weeks to collect baseline GSM values. Dorsal hemisection injuries and intra-spinal cord injections of AAV-rg/Cre were done when mice were 17 weeks of age. Mice received 4 bilateral injections, 2 above and 2 below the injury (total of 12E9 genome copies (GC) per mouse, 0.3 µl/injection in the same surgery as a dorsal hemisection injury at C5 (hemisections were done first). Of those, 3 PTEN^f/f^;Rosa^tdTomato^ mice and 1 Rosa^tdTomato^ mouse died 1-2 days post-surgery. Thus after attrition, starting animal numbers were 12 PTEN^f/f^;Rosa^tdTomato^ mice (n=8 male; n=4 female) and 13 Rosa^tdTomato^ mice (n=4 male; n=9 female).

Starting 4 weeks post-injury, vivarium staff began reporting wounds behind the ears of many mice. Details on these are described in the Results, as well as our determination of the cause (incessant scratching). Despite treatment, wounds increased in severity to the point that some mice were euthanized. Some mice also began to exhibit rigid forward extension of the hindlimbs, which we term dystonia. Because of the increasing severity of the pathophysiologies, the study was terminated and mice were euthanized 9 weeks post AAV-rg/Cre injections plus SCI.

Importantly, mice were group housed (up to 4 mice/cage), treatment groups were not separated, and both vivarium staff and individuals doing GSM testing were blind to genotype. As a result, we did not detect the fact that pathophysiologies were occurring only in PTEN^f/f^;Rosa^tdTomato^ mice with SCI and AAV-rg/Cre until the study was terminated and blinding was removed. Overall, 9/13 PTEN^f/f^;Rosa^tdTomato^ mice with SCI and AAV-rg/Cre exhibited wounds and scratching and some hindlimb dystonia.

To begin to define causes of the pathophysiologies, we carried out follow-up studies. In Study 2, we reduced AAV-rg doses (6E9 GC/animal) to determine if this was related to the occurrence of pathophysiologies. Injections were made rostral to the injury rather than both rostral and caudal as in Study 1. Study 2 involved a total of 26 PTEN^f/f^;Rosa^tdTomato^ mice that were 11 weeks old at the start of the study. Mice were handled for 2 weeks; GSM testing began when mice were 13 weeks old and continued for 2 weeks to collect baseline GSM values. At 15 weeks of age, mice received dorsal hemisection injuries and bilateral intra-spinal cord injections of AAV above the injury. 16 mice (n=9 male; n=7 female) received 6E9 GC per mouse of AAV-rg/Cre. 10 mice (n=6 male; n=4 female) received 6E9 GC of the control vector AAV-rg/GFP (0.3 µl/injection). Of those, 9 mice injected with AAV-rg/Cre and 3 mice injected with AAV-rg/GFP died on the first 1-2 days post-surgery. Thus after attrition, starting animal numbers were a total of 18 PTEN^f/f^;Rosa^tdTomato^ mice; 11 mice (n=7 male; n=4 female) injected with AAV-rg/Cre and 7 mice (n=4 male; n=3 female) injected with AAV-rg/GFP.

Two PTEN^f/f^;Rosa^tdTomato^ mice injected with AAV-rg/Cre began to exhibit pathophysiologies 5 weeks post injections/SCI, although pathophysiologies were not as severe as seen with higher doses of AAV-rg/Cre in Study 1, and we were able to conduct the experiment to its completion. All mice were perfused 16 weeks post AAV-injection/ SCI.

In Study 3 we further reduced AAV-rg dose to 3E9 GC/animal. Here, injections were done rostral to the injury, but only on one side. Onset and nature of pathophysiologies was systematically monitored. Study 3 involved a total of 30 PTEN^f/f^;Rosa^tdTomato^ mice (n=14 male; n=16 female); controls were 16 Rosa^tdTomato^ mice (n=8 male; n=8 female). Mice were 25 weeks old at the start of the study and were handled for 2 weeks. GSM testing began when mice were 27 weeks old and baseline values were collected for 9 weeks. At 36 weeks of age, mice received dorsal hemisection injuries and unilateral intra-spinal cord injections of AAV-rg/Cre above the lesion (0.3 µl/injection). 5 PTEN^f/f^;Rosa^tdTomato^ mice and 1 Rosa^tdTomato^ mouse died 1-2 days post-surgery, thus after attrition, total starting animal numbers were 25 PTEN^f/f^;Rosa^tdTomato^ mice (n=10 male; n=15 female) and 15 Rosa^tdTomato^ mice (n=8 male; n=7 female).

Despite lower dose of AAV-rg/Cre, mice in Study 3 still exhibited pathophysiologies (detailed in Results). However, only one mouse was euthanized due to the severity of the pathophysiology, so remaining mice were tested for the planned duration of the study. Mice were perfused 12 weeks post AAV injections/SCI.

Study 4 assessed consequences of shRNA-based knockdown of PTEN in conjunction with SCI. 15 Rosa^tdTomato^ mice (n=7 male; n=8 female) received AAV-rg/shPTEN/GFP, while controls were 15 Rosa^tdTomato^ mice that received AAVrg/GFP (n=8 male; n=7 female). Mice were 25 weeks old at the start of the study, were handled for 2 weeks, and then baseline GSM values were collected for 9 weeks. At 34 weeks of age, mice received dorsal hemisection injuries and unilateral intra-spinal cord injections of 3E9 GC per mouse of AAV-rg above the lesion (0.3 µl/injection). 1 mouse injected with AAV-rg/shPTEN/GFP died within the first 2 days post-surgery, and 1 mouse injected with AAV-rg/GFP died 1 day after SCI. Thus, after attrition, starting animal numbers were 14 (n=6 male; n=8 female) for Rosa^tdTomato^ mice injected with AAV-rg/shPTEN/GFP and 14 (n=8 males; n=6 females) for the group injected with AAV-rg/GFP.

Of the mice that received AAV-rg/shPTEN/GFP, one male mouse exhibited hindlimb dystonia 1 week post lesion/injection, and excessive scratching. This mouse was euthanized at this time point (1 week post lesion/injection) due to severity of scratching wounds. One male mouse developed shoulder wounds due to scratching 2.5 weeks post lesion/injection and 6 more (n=3 males; n=3 females) began to exhibit pathophysiologies 5 weeks post lesion/injection. The scratching wounds were treated and did not become severe enough to require euthanasia. All other mice continued in the study and were perfused 12 weeks post lesion/injection.

Study 5 tested whether pathophysiologies were due to deletion of PTEN in local spinal cord circuitry in the context of a spinal cord injury. For this, mice received AAV2/Cre rather than AAV-rg/Cre at the time of SCI to delete PTEN in the spinal cord around the injection site without retrograde transduction of cells of origin of spinal pathways in the brain. 15 PTEN^f/f^;Rosa^tdTomato^ (n=6 male; n=9 female) and 13 Rosa^tdTomato^ mice (n=8 male; n=5 female) were 12 weeks old at the start of the study, were handled for 2 weeks and baseline GSM values were collected for 2 weeks. At 16 weeks of age, mice received dorsal hemisection injuries and 2 bilateral intra-spinal cord injections of AAV2/Cre above the injury (0.3 µl/injection). Of these, 1 PTEN^f/f^;Rosa^tdTomato^ mouse and 1 Rosa^tdTomato^ mouse died on the first 1-2 days post-surgery; thus after attrition, total starting animal numbers were 14 PTEN^f/f^;Rosa^tdTomato^ mice (n=5 male; n=9 female) and 12 Rosa^tdTomato^ mice (n=8 male; n=4 female). No pathophysiologies were seen, and these mice were perfused 8 weeks post AAV2/Cre injections plus SCI.

Several of our previous anatomical studies involved injections of AAV-rg/Cre in PTEN^f/f^;Rosa^tdTomato^ mice and Rosa^tdTomato^ mice without injuries (Metcalfe et al., 2022). Thus, we respectively reviewed husbandry reports and survival statistics to determine whether pathophysiologies occurred with AAV-rg/Cre injections in PTEN^f/f^;Rosa^tdTomato^ mice in the absence of spinal cord injury.

The retrospective review included n=12 PTEN^f/f^;Rosa^tdTomato^ mice (n=8 males were between 14-24 weeks old; n=3 females were between 16-24 weeks old). These mice received two bilateral intra-spinal cord injections of AAV-rg/Cre (6E9 GC/mouse) at C5 as in study 2 here. No animals died during injection procedures. Mice were allowed to survive for different time periods and were killed for anatomical studies. Two males and 1 female mouse were killed 1 month post injection; 4 male mice were killed 4 months post injection; 1 male mouse was killed 7 months post injection and 2 male and 3 females were killed 12 months post injection. As detailed in the Results, none of these mice exhibited pathophysiologies.

Details on animal use (genotypes, ages at the time of injection, injection location, survival time) are summarized in Table 1.

### 2.5. Grip strength meter (GSM)

The GSM measures grip strength (the maximal peak force in grams) as a mouse is allowed to grip a bar and then is gently pulled away. To test grip strength of one paw, the opposite forepaw is taped with non-stick surgery tape (Danilov and Steward, 2015). For testing, mice were held by the tail and suspended 2-3 inches above the plate of the GSM. Mice were gradually brought toward the bar until they reached out and gripped the bar. Mice were allowed to briefly adjust their paw on the bar to establish grip, then the mice were gently pulled away until the grip was released. The GSM records the maximal force at the time grip was released. This procedure was repeated 4 times in each testing session, recording grip strength separately for each paw and then averaging across the session.

For each study, mice were pre-handled for 2 weeks and then pre-trained on the GSM as stated for each study. Mice typically exhibit a stronger grip early in training and then gripping force decreases to a plateau that is maintained over time; consequently, only the last 5 days of pre-injury values are included in the graphs. After GSM training, mice received dorsal hemisections and were then tested post injury on the GSM 3x per week as stated for each study.

### 2.6. Tissue preparation

Mice were transcardially perfused at survival times indicated in Table 1 with 4% paraformaldehyde in 0.1M phosphate buffer (4% PFA). Brains, and spinal cords with attached dorsal root ganglia were dissected, post-fixed in 4% PFA overnight, then immersed in 27% sucrose for cryoprotection overnight, frozen in TissueTek OCT (VWR International) and stored at −20°C until they were sectioned with a cryostat.

### 2.7. Immunostaining to assess AAV transduction

Brains were sectioned at 30 µm in the coronal plane collecting sets of sections at 240 µm intervals to generate 10 sets of regularly-spaced sections. One set of the sections was mounted in serial order for visualization of native tdT fluorescence to assess AAV transduction. Other sets of sections were immunostained for tdT (rabbit anti-red fluorescent protein, Rockland Immunochemicals Inc., #600-401-379, RRID:AB_2209751). To assess deletion/knockdown, one set of sections was immunostained for PTEN (rabbit anti-PTEN Cell Signaling Technology #9188L, RRID:AB_2253290). Biotin-conjugated secondary antibodies were used at 1:250 dilution and stained with tyramide-Cy3 (for PTEN).

### 2.8. Quantification of PTEN-deleted cortical motoneurons (CMNs)

One set of sections at 240 µm intervals was prepared for PTEN immunofluorescence (IF) to detect PTEN-deleted cortical motoneurons (CMNs). PTEN-deleted neurons appear as unlabeled “ghost-cells” against the background fluorescence over the neuropil. In PTEN^f/f^;Rosa^tdTomato^ mice, neurons transduced by AAV-rg/Cre express tdTomato (tdT), and native tdT fluorescence is visible in sections prepared for PTEN immunofluorescence allowing co-imaging, which confirmed that tdT-positive neurons also negative for PTEN (Fig. 1).

**Figure 1.**
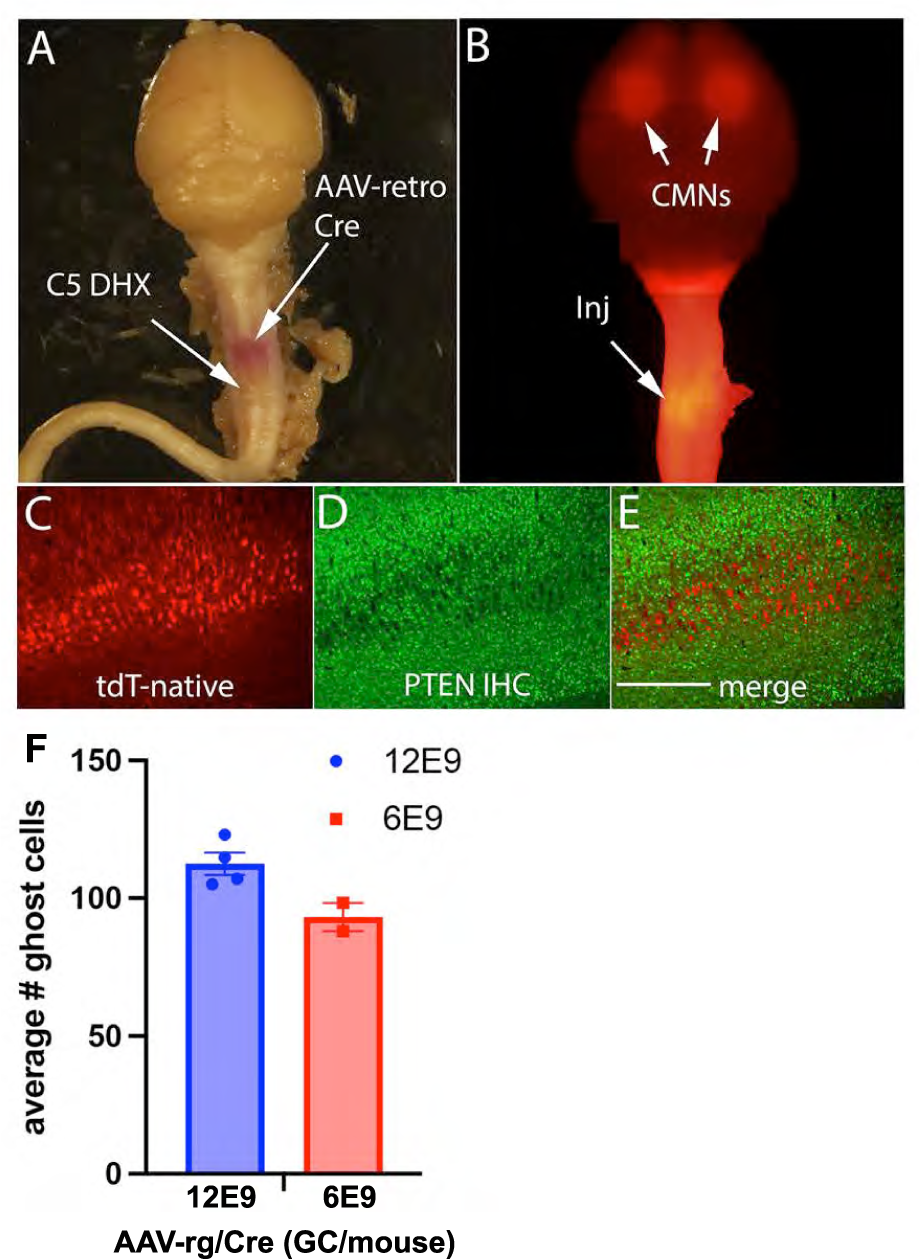
Retrograde transfection of cortical motoneurons (CMNs) with injections of retro-AAV/Cre into the spinal cord of PTEN^f/f^: Rosa^tdTomato^ mice. A) Intact brain and attached spinal cord. tdTomato is visible at the site of injection in the cervical spinal cord as well as the site of injury (arrows) B) Epifluorescence images of the same brain and spinal cord showing oval areas of tdT fluorescence marking clouds of CMNs expressing tdT and site of injection in thecervical spinal cord (arrows). C-E) Coronal sections through the sensorimotor cortex showing large numbers of native fluorescent tdT-positive pyramidal neurons in layer V (C) that were completely negative for PTEN protein (D). Co-imaging for tdT revealed that tdT-positive neurons are negative for PTEN protein (E). F) Different doses of AAV-retro/Cre revealed an average of 112.5 ghost cells/section at the 12E9 GC dose, 93.16 ghost cells per section at the 6E9 GC/mouse dose, and 42.7 ghost cells at the 3E9 dose. One-way ANOVA revealed significant differences between GC doses [F(64.96) DF 2, 6, p<0.0001]. Post-hoc Tukey’s multiple comparisons test revealed significant differences between high (12E9 GC/mouse) and low GC doses (3E9 GC/mouse) (p<0.0001, 95% C.I. = [46.6, 81.27] and between intermediate (6E9 GC/mouse) and low GC doses (3E9 GC/mouse) (p = 0.0014, 95% C.I. = [23.88, 65.32]). There were no significant differences between high (12E9 GC/mouse) and intermediate GC doses (6E9 GC/ml) (p = 0.0532, 95% C.I. = [-0.32, 39]).

Ten sections spanning 2.4 mm along the A-P axis were used for quantitative analyses of numbers of PTEN-deleted neurons (ghost cells). For animals in the high dose group (12E9 GC/mouse), n=4 mice were chosen taking the first three in the study list that were not used for analyses for other studies (Metcalfe et al., 2022). For mid doses (6E9 GC/mouse) n=2 mice were chosen from the top and the bottom of the study list and that survived until the end of the study. The area of each section containing PTEN-deleted neurons was imaged at 10X. Images of cortex containing ghost cells were obtained by setting edge points encompassing the totality of layer V in that section at 10x magnification using the Keyence microscope “Image Stitch” program. Keyence analysis software stitched the individual frames together and measurements were then exported to ImageJ (Schneider et al., 2012) to determine the average number of PTEN-negative neurons (ghost cells) per section and then average number of ghost-cells per mouse. Measurements were then exported to Excel (Microsoft) and an average of ghost-cells per mouse for each dose was determined.

### 2.9. Statistical Analyses

Prism software was used for all statistical analyses. For statistical analysis of ghost cells numbers, the analysis was a one-way ANOVA with a post hoc Tukey’s multiple comparisons test to determine significant differences among groups. For GSM values analyses, the primary statistical analysis was by a two-way repeated measures ANOVA for each separate study with variables of “groups” and “time post-injury”. A mixed effects model analysis was used when values were missing for some timepoints, particularly when animals were euthanized due to health concerns. We also calculated *percent recovery in each group* by comparing average pre-injury GSM values vs. peak post injury values. Statistical analyses then compared groups (PTEN deleted vs. control) with values being the percent recovery for each animal.

## 3. RESULTS

### 3.1 Intra-spinal cord injections of AAV-rg/Cre in PTEN^f/f^;Rosa^tdTomato^ mice transduce corticomotor neurons to induce tdT expression and delete PTEN

The present study takes advantage of a model using double-transgenic mice (PTEN^f/f^;Rosa^tdTomato^) and AAV-rg/Cre, which allows remote transduction of neurons via retrograde transport. We have previously shown that intra-spinal cord injections of AAV-rg/Cre in PTEN^f/f^;Rosa^tdTomato^ mice led to selective transduction of thousands of neurons in layer V of the sensorimotor cortex as revealed by induction of tdTomato (tdT) expression. These are the cells of origin of the corticospinal tract (CST), termed “corticomotor neurons” (CMNs). Cells of origin of other descending spinal pathways were also transduced, including neurons in the brainstem reticular formation, red nucleus, and hypothalamus (Metcalfe et al., 2022). Studies from other groups have revealed that AAV-rg also transduces the cells of origin of some but not all intra-spinal circuits; for example, neurons of the propriospinal pathways and serotoninergic neurons in the midline raphe nuclei of the medulla are not efficiently transduced (Wang et al., 2018). Finally, some populations of neurons in dorsal root ganglia near the injection are also transduced (Metcalfe et al., 2022).

Results of the present studies were consistent with previous data. In Study 1, PTEN^f/f^;Rosa^tdTomato^ mice received doses of AAV-rg/Cre that were 2-fold higher than in our previous study (total of 12E9 GC/mouse) at the time of a dorsal hemisection. This was delivered in 4 injections, 2 above and 2 below the injury. TdT was visible in the spinal cord at the injection site in intact spinal cords (Fig. 1A), and epi-fluorescence illumination revealed oval areas of tdT fluorescence in the cortex marking clouds of CMNs expressing tdT (Fig. 1B). In coronal sections through the sensorimotor cortex, large numbers of tdT-positive pyramidal neurons were evident in layer V without immunostaining (native fluorescence, Fig. 1C).

Confirming previous findings, immunostaining for PTEN in PTEN^f/f^;Rosa^tdTomato^ mice injected with AAV-rg/Cre revealed large numbers of CMNs that were completely negative for PTEN protein, appearing as “ghost cells” (Fig. 1D, same section as Fig. 1C). Co-imaging for tdT revealed that tdT-positive neurons are negative for PTEN protein (No double labeling, which would be represented in yellow, Fig. 1E).

Different doses of AAV-rg/Cre were delivered in studies 1, 2, and 3 (12E9 GC/mouse delivered in two injections above the lesion in study 1, 6E9 GC/mouse delivered in two injections above the lesion in study 2, and 3E9 GC/mouse in one unilateral injection in Study 3). Counts revealed an average of 112.5 ghost cells/section throughout the sensorimotor cortex at the 12E9 GC dose and 93.16 ghost cells per section at the 6E9 GC/mouse dose (Fig. 1F). Mice in study 3 received intra-cortical BDA injections prior to sacrifice, which cause some cortical damage, making the tissue unsuitable for counts of ghost cells. Two-tailed t-test analysis revealed significant differences between high (12E9 GC/mouse) and intermediate GC doses (6E9 GC/ml) (t(4.00) = 2.790, p = 0.0454). The likely explanation for the small difference between the high and intermediate GC doses in average numbers of PTEN-negative cells/section is that 2 of the injections at the high dose were below the injury. With complete dorsal hemisections, injections below the injury would not be expected to transduce many CMNs because the dorsal and dorso-lateral CST is completely transected.

### 3.2. Enhanced recovery of forelimb motor function after SCI with intra-spinal cord injections of AAV-rg/Cre

#### 3.2.1. Study 1

The rationale of Study 1 was to test AAV-rg as a platform to delete PTEN in multiple descending and ascending pathways that would be transected by a dorsal hemisection injury. For this, mice received dorsal hemisection injuries at C5 and bilateral injections of AAV-rg/Cre 1mm rostral and 1mm caudal to the injury (total of 12E9 GC/mouse). Controls were Rosa^tdTomato^ mice that received comparable injections. Forelimb motor function was assessed over time using a GSM by an individual blinded to treatment group.

Following C5 lesions and AAV-rg/Cre intra-spinal cord injections, GSM values dropped after injury to 5.5% of pre-injury values for Rosa^tdTomato^ mice and 26.9% of pre-injury values for PTEN^f/f^;Rosa^tdTomato^ mice (Fig. 2A). For the control Rosa^tdTomato^ mice GSM values recovered to a plateau at around 14 days (36% of pre-injury control). In contrast, values for PTEN^f/f^;Rosa^tdTomato^ mice increased progressively reaching a plateau at 25 days of 84% of pre-injury control values. At all post-injury time points, average values for the PTEN^f/f^;Rosa^tdTomato^ mice (PTEN-deleted) were higher than Rosa^tdTomato^ controls. Two-way repeated measures ANOVA revealed significant differences over time [F(15.13) DF 18, 414, p<0.0001] and between genotypes [F(6.41) = DF 1,23, p=0.018].

**Figure 2.**
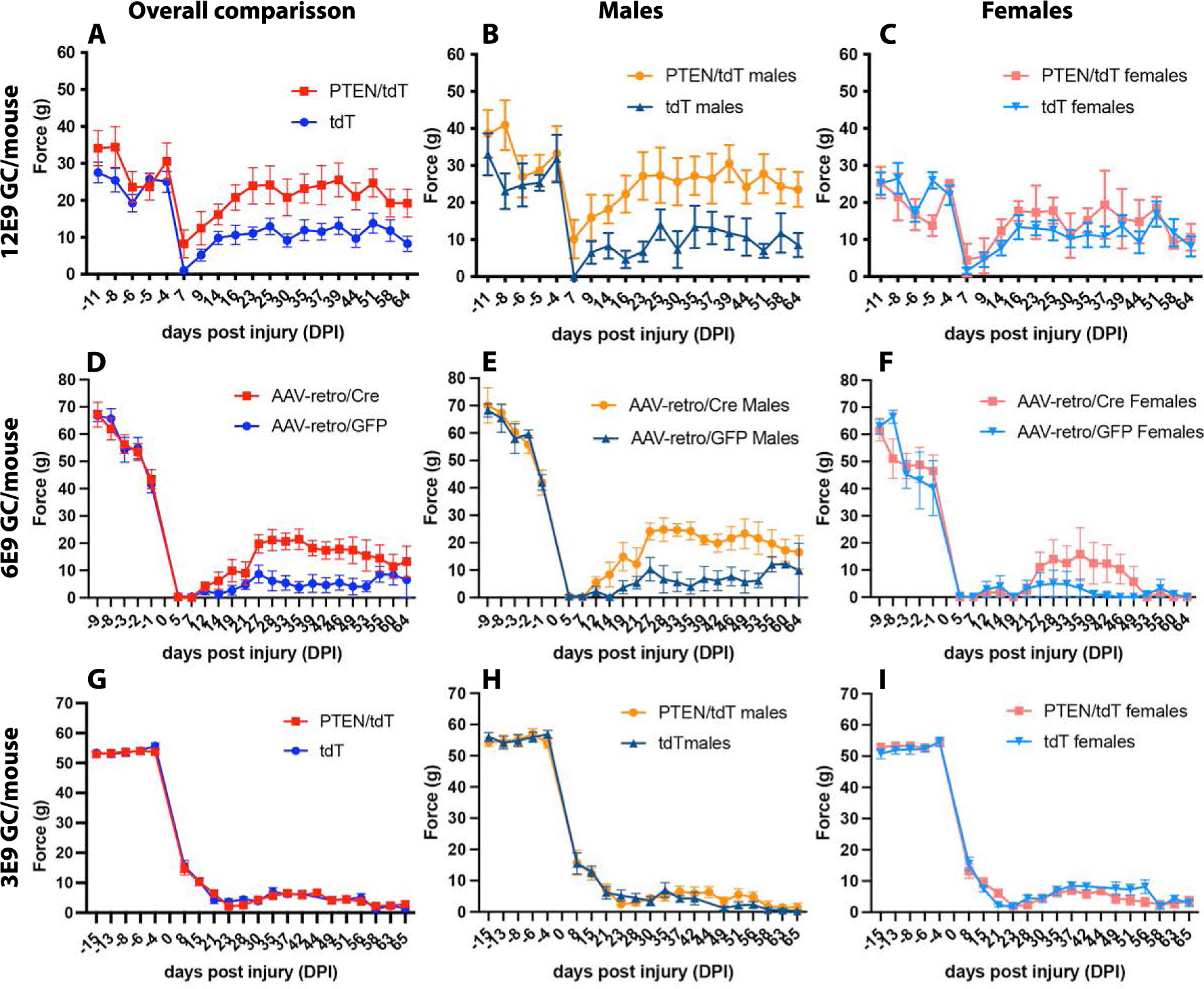
Genetic deletion of PTEN promotes transient functional recovery followed by functional deterioration. Forepaw gripping function as measured by grip strength meter (GSM). A) Rosa^tdTomato^ and PTEN^f/f^;Rosa^tdTomato^ mice were injected with 12E9 GC/mouse of AAV-retro/Cre at the same time as a cervical dorsal hemisection. Two-way repeated measures ANOVA revealed significant differences over time [F(15.13) DF 18, 414, p<0.0001] and between genotypes [F(6.41) = DF 1,23, p=0.018]. B-C) Stratification of GSM data by sex revealed no sex differences in the values of the Rosa^tdTomato^ control mice [F(0.63) DF 1,11, p=0.442], whereas there were substantial sex differences in PTEN^f/f^;Rosa^tdTomato^ mice (PTEN-deleted) [F(4.97) DF 1,10, p=0.05]. D) PTEN^f/f^;Rosa^tdTomato^ mice were injected with 6E9 GC/mouse of AAV-retro/Cre or AAV-retro/GFP at the same time as a cervical dorsal hemisection. A mixed effects model analysis revealed significant differences over time [F(121.1) DF 5.07, 65.4, p<0.0001] and between genotypes [F(5.47) = DF 1, 14, p=0.035]. E-F) Stratification of GSM data by sex revealed no sex differences in the values of the AAV-retro/GFP control mice (PTEN intact) [F(1.63) DF 1,5, p=0.26] but significant sex differences for the PTEN-deleted mice injected with AAV-retro/Cre [F(8.39) DF 1,7, p=0.023]. G) Rosa^tdTomato^ and PTEN^f/f^;Rosa^tdTomato^ mice were injected with 3E9 GC/mouse of AAV-retro/Cre at the same time as a cervical dorsal hemisection. A mixed effects model analysis revealed significant differences over time [F(823.0) DF 4.56, 174.6, p<0.0.001] but no significant differences between genotypes [F(0.018) = DF 1, 39, p=0.8944]. H-I) A mixed model ANOVA revealed no significant sex differences for the Rosa^tdTomato^ mice [F(0.92) DF 1,13, p=0.35] nor for the PTEN^f/f^;Rosa^tdTomato^ mice [F(2.29) DF 1,23, p=0.63].

##### 3.2.1.a. Comparisons by sex

Stratification of GSM data by sex (Fig. 2B &C) revealed no sex differences in the values of the Rosa^tdTomato^ control mice. However, there were substantial sex differences in PTEN^f/f^;Rosa^tdTomato^ mice (PTEN-deleted). Specifically, enhanced recovery of GSM values was seen primarily in male mice. GSM values for female PTEN^f/f^;Rosa^tdTomato^ mice (PTEN-deleted) were 77.3% of pre-injury control at the plateau vs. 35.6% for the Rosa^tdTomato^ female control mice. In contrast, GSM values for male PTEN^f/f^;Rosa^tdTomato^ mice were 95.6% of pre-injury control at the plateau vs. 27.1% for the Rosa^tdTomato^ male control mice. Two-way repeated measures ANOVA revealed no significant sex differences for the Rosa^tdTomato^ mice [F(0.63) DF 1,11, p=0.442] but significant sex differences for the PTEN^f/f^;Rosa^tdTomato^ mice [F(4.97) DF 1,10, p=0.05]. Thus, the values for the male PTEN^f/f^;Rosa^tdTomato^ mice largely accounted for the overall difference in the combined values reported above.

##### 3.2.1.b. Late-developing pathophysiologies

Mice were tested with GSM once/week and after the initial post-injury period; mice were not individually examined every day by research staff but were monitored routinely by vivarium staff (routine animal husbandry). At 4 weeks post-injury, vivarium staff reported a mouse with severe skin wounds behind the ears and on the neck. These wounds were initially attributed to “over-grooming”. After careful observation, we determined that the mouse was repeatedly scratching the area of the wound with its hindlimbs. Wounds on this mouse were severe so the mouse was euthanized. At 5 weeks, another mouse was reported as having wounds diagnosed as “ulcerative dermatitis”. Wounds were treated with Silvadeen and toenails were trimmed. Similar wounds began to appear in other mice that were attributed to various causes; in each case, wounds were treated with Silvadeen and toenails were trimmed. At 6 weeks, 3 more mice exhibited wounds that were diagnosed as “skin lesions” and “ulcerative dermatitis”. At 7 weeks, 2 more mice exhibited wounds that were diagnosed as “fight wounds and ulcerative dermatitis”. Finally, at 9 weeks, 2 more mice exhibited wounds diagnosed as “fight wounds” in one case and “ulcerative dermatitis” in the other. Some mice also began to exhibit rigid forward extension of the hindlimbs, which we term dystonia. Figure 3 illustrates the nature of these late-onset pathophysiologies.

**Figure 3.**
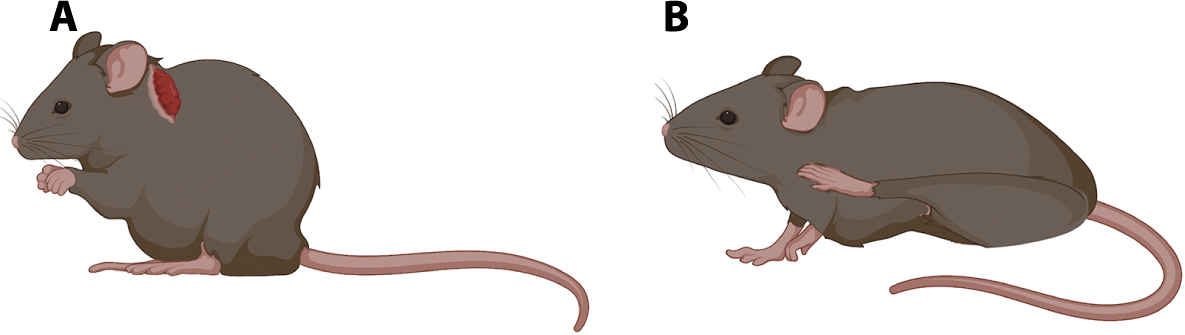
Pathophysiology with AAV-rg mediated PTEN deletion and SCI. At 2-3 months post-injury animals develop excessive scratching and hindlimb dystonia. PTEN^f/f^;Rosa^tdTomato^ injected with AAV-rg/Cre show severe scratching (A) and/or hindlimb dystonia (B).

Overall, 9/13 PTEN^f/f^;Rosa^tdTomato^ mice with SCI and AAV-rg/Cre exhibited wounds and scratching. We had originally planned to inject BDA to trace CST axons, but because of the continued incessant scratching and increasing severity of the wounds, we terminated the experiment and all remaining mice were euthanized and perfused at 64 days post-AAV-rg/Cre injections. When mice were terminated, we did not note the sex of mice that had exhibited pathophysiologies, and whether there were sex differences in the incidence or severity of wounds.

#### 3.2.2. Study 2

To begin to define causes and mechanisms of the pathophysiologies that appeared in study 1, in study 2, we made 2 adjustments to the dosing. First, the overall dose was reduced to 6E9 GC/mouse to determine whether this dose could promote forelimb recovery without occurrence of pathophysiologies. Second, because some mice exhibited rigid forward extension of the hindlimb, we wondered whether this would be reduced or eliminated if AAV-rg/Cre injections were only rostral to the injury. Also, for controls, genotype was held constant; all mice were PTEN^f/f^;Rosa^tdTomato^ but the control group was injected with AAV-rg/GFP rather than AAV-rg/Cre.

One week following C5 lesions and AAV-rg injections, GSM values on the first testing day were 1% of pre-injury values for mice injected with AAV-rg/Cre and 1.7% of pre-injury values for mice injected with AAV-rg/GFP. For the control mice injected with AAV-rg/GFP, GSM values recovered progressively to a plateau at around 25 days post-injury (9.1% of pre-injury control). In contrast, values for PTEN-deleted mice increased progressively reaching a plateau at 25 days 49.3% of pre-injury control values (Fig. 2D). A mixed effects model analysis revealed significant differences over time [F(121.1) DF 5.07, 65.4, p<0.0001] and between genotypes [F(5.47) = DF 1, 14, p=0.035].

Of note, post-injury values for controls were lower in this study (9% of pre-injury) in comparison to study 1 (36% of pre-injury). One possible explanation is that the control group is differs between studies (Rosa^tdTomato^ in study 1 vs. PTEN^f/f^;Rosa^tdTomato^ that received AAV-rg/GFP in study 2). Differences in lesion extent are possible but less likely because the same surgeon did the dorsal hemisections for all studies. The other possibility is that different researchers tested mice by GSM in the two studies. Because of the differences in GSM testers between study 1 and 2 we defined more strict criteria for GSM scoring to improve inter-rater reliability for subsequent studies.

##### 3.2.2.a. Comparisons by sex

Stratification of GSM data by sex (Fig. 2E &F) revealed no sex differences in the values of the AAV-rg/GFP control mice (PTEN intact). However, as in Study 1, there were substantial sex differences in PTEN^f/f^;Rosa^tdTomato^ mice injected with AAV-rg/Cre (PTEN-deleted). Again, enhanced recovery of GSM values was greater in male mice. GSM values for female PTEN-deleted were 30.25% of pre-injury control at the plateau vs. 7.86% for the PTEN-intact female control mice. In contrast, GSM values for male PTEN-deleted mice were 61.1% of pre-injury control at the plateau vs. 11.8% for the PTEN-intact male control mice. A mixed effects model analysis revealed no significant sex differences for control mice injected with AAV-rg/GFP [F(1.63) DF 1,5, p=0.26] but significant sex differences for the PTEN-deleted mice injected with AAV-rg/Cre [F(8.39) DF 1,7, p=0.023]. Thus, as in study 1, the values for the male PTEN^f/f^;Rosa^tdTomato^ mice largely accounted for the overall difference in the combined values.

##### 3.2.2.b. Late-developing pathophysiologies

Although mice that received AAV-rg/Cre exhibited higher GSM scores throughout the testing period, pathophysiologies developed at 5 weeks post injection/SCI in n=2 PTEN^f/f^;Rosa^tdTomato^ mice (n=1 male; n=1 female). The male mouse was reported as having hindlimb dystonia plus autophagia and the female exhibited scratching wounds on shoulders. These mice were euthanized because of the severity of the wounds. Behavioral testing (GSM) continued for 64 days for the remaining mice [7 PTEN^f/f^;Rosa^tdTomato^ mice injected with AAV-rg/Cre (n=5 male; n=2 female) and 7 injected with AAV-rg/GFP (n=4 male; n=3 female)]. We did not record which animal numbers exhibited pathophysiologies, so in this study we were unable to determine if there were sex differences in the incidence of wounds. None of the controls (PTEN^f/f^;Rosa^tdTomato^ mice injected with AAV-rg/GFP, PTEN-intact) exhibited excessive scratching, wounds, or hindlimb dystonia at any time. Mice were perfused 16 weeks post injection/SCI.

#### 3.2.3. Study 3

In study 3 we further decreased AAV-rg/Cre doses (3E9 GC/animal). PTEN^f/f^;Rosa^tdTomato^ mice received 1 unilateral injection above the level of the dorsal hemisection at approximately C4. Controls were Rosa^tdTomato^ mice that received comparable injections. Forelimb motor function was assessed over time using a GSM by an individual blinded to treatment group.

Following C5 lesions and AAV-rg/Cre injections, (one week post-injury), GSM values decreased to 4.1% of pre-injury values for Rosa^tdTomato^ mice and 6.4% of pre-injury values for PTEN^f/f^;Rosa^tdTomato^ mice. For the control Rosa^tdTomato^ mice GSM values recovered to a plateau at around 25 days post-injury (6.8% of pre-injury control). Values for PTEN^f/f^;Rosa^tdTomato^ mice were slightly higher than control mice, reaching a plateau at 25 days 12.4% of pre-injury control values (Fig. 2G). A mixed effects model analysis revealed significant differences over time [F(823.0) DF 4.56, 174.6, p<0.0.001] but no significant differences between genotypes [F(0.018) = DF 1, 39, p=0.8944]. Since injections were unilateral, we analyzed GSM values per paw (right and left). A mixed effects model analysis revealed significant differences over time on the left [F(470.2) DF 5.14, 197.9, p<0.0.001] and the right paw [F(606.7) DF 6.79, 261.5, p<0.0.001] but no significant differences between genotypes on the left [F(0.545) DF 1, 40, p=0.465] or the right paw [F(0.00059) = DF 1, 40, p=0.981] (Fig. 4).

**Figure 4.**
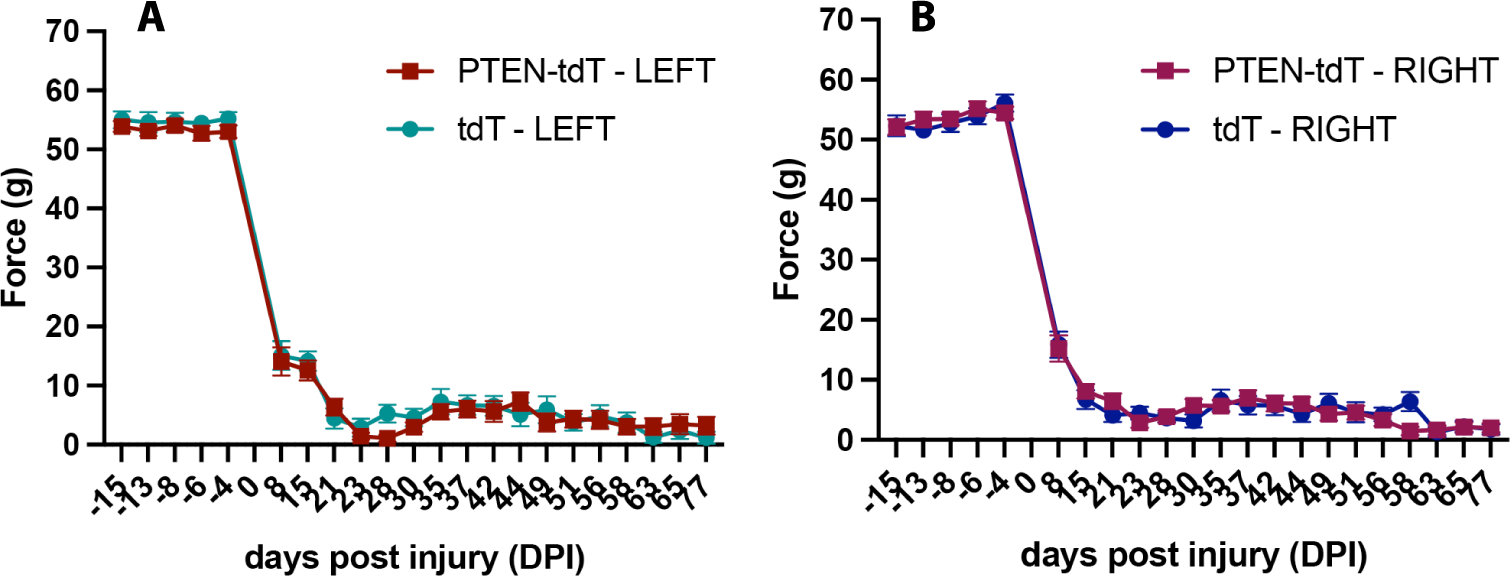
Unilateral low doses of AAV-rg/Cre show no difference in paw performance. Since injections were unilateral, we analyzed GSM values per paw (right and left). A mixed effects model analysis revealed significant differences over time on the left [F(470.2) DF 5.14, 197.9, p<0.0.001] and the right paw [F(606.7) DF 6.79, 261.5, p<0.0.001] but no significant differences between genotypes on the left [F(0.545) DF 1, 40, p=0.465] or the right paw [F(0.00059) = DF 1, 40, p=0.981] (Fig. 4).

##### 3.2.3.a. Comparisons by sex

Stratification of GSM data by sex (Fig. 2H &I) revealed no sex differences in the values of the Rosa^tdTomato^ control mice or PTEN^f/f^;Rosa^tdTomato^ mice (PTEN-deleted). Specifically, GSM values for female PTEN^f/f^;Rosa^tdTomato^ mice (PTEN-deleted) were 12.6% of pre-injury control at the plateau vs. 15.6% for the Rosa^tdTomato^ female control mice. GSM values for male PTEN^f/f^;Rosa^tdTomato^ mice were 13% of pre-injury control at the plateau vs. 7.7% for the Rosa^tdTomato^ male control mice. A mixed model ANOVA revealed no significant sex differences for the Rosa^tdTomato^ mice [F(0.92) DF 1,13, p=0.35] nor for the PTEN^f/f^;Rosa^tdTomato^ mice [F(2.29) DF 1,23, p=0.63]. Overall neither group showed improvement in recovery of function after SCI plus AAV-rg/Cre injection.

Again, post-injury values for controls were lower in this study (6.8% of pre-injury) in comparison to study 1 (36% of pre-injury). In this case, the control group is the same (Rosa^tdTomato^). Differences in lesion extent are possible but less likely because the same surgeon did the dorsal hemisections for all studies. The possible explanation that remains is that the GSM tester in study 1 was different than in studies 2 and 3.

##### 3.2.3.b. Late-developing pathophysiologies

In study 3, despite lowering the dose of AAV-rg/Cre to the extent that functional recovery due to PTEN deletion was lost, PTEN^f/f^;Rosa^tdTomato^ mice (PTEN-deleted) mice still developed late occurring pathophysiologies. The first incidence of pathophysiology was reported 3 weeks post AAV-rg/Cre injection and SCI in n=4 mice (all males). N=3 of those showed injuries behind their ears due to chronic scratching and n=1 exhibited hindlimb dystonia. Of those, 1 of the animals that were scratching subsequently developed hindlimb dystonia at 7 weeks post AAV-rg/Cre injections plus SCI. This mouse was euthanized due to severity of injuries at this time point. New cases of pathophysiologies appeared at 7 weeks post AAV-rg/Cre injections plus SCI in n=9 mice (n=5 males; n=4 females). Of these, n=4 showed only scratching behind the ears, n=1 only hindlimb dystonia and n=4 exhibited both scratching and hindlimb dystonia. Injuries were not as severe as in study 1, so no other animals were euthanized due to severity of pathophysiologies. Thus out of n=26 PTEN^f/f^;Rosa^tdTomato^ mice (n=10 males; n=16 females), 8/10 males and 5/16 females developed pathophysiologies starting 3 weeks post AAV-rg/Cre injections plus SCI, These results suggest that the incidence of pathophysiologies in males is higher compared to females. Animals were perfused 12 weeks post AAV-rg/Cre injections plus SCI.

#### 3.2.4. Study 4: AAV-rg/shRNA-based knockdown of PTEN

Study 4 assessed whether PTEN knockdown via AAV-rg/shPTEN would enhance motor recovery and/or lead to pathophysiologies. Rosa^tdTomato^ mice received unilateral intraspinal cord injections of AAV-rg/shPTEN/GFP above the injury (3E9 GC/mouse). Controls were Rosa^tdTomato^ mice that received similar injections of AAV-rg/Cre. Forelimb motor function was assessed over time using a GSM by an individual blinded to treatment group.

Following C5 lesions and AAV-rg injections, GSM values were 8% of pre-injury values at 21 days post-injury for mice injected with AAV-rg/Cre (PTEN-intact controls). Earlier time points are not shown because of errors in scoring. Values for mice that received AAV-rg/shPTEN were 5% of pre-injury control. For the control PTEN-intact mice GSM values recovered to a plateau at around 37 days post-injury of 11.8% of pre-injury control, whereas values for Rosa^tdTomato^ mice that received AAV-rg/shPTEN were 6.7% of pre-injury control. A mixed effects model analysis revealed significant differences over time [F(535.8) DF 25, 375, p<0.0.001] and but no significant differences between PTEN-deleted and PTEN-intact groups [F(0.8753) = DF 1, 15, p=0.4301] (Fig. 5A).

**Figure 5.**
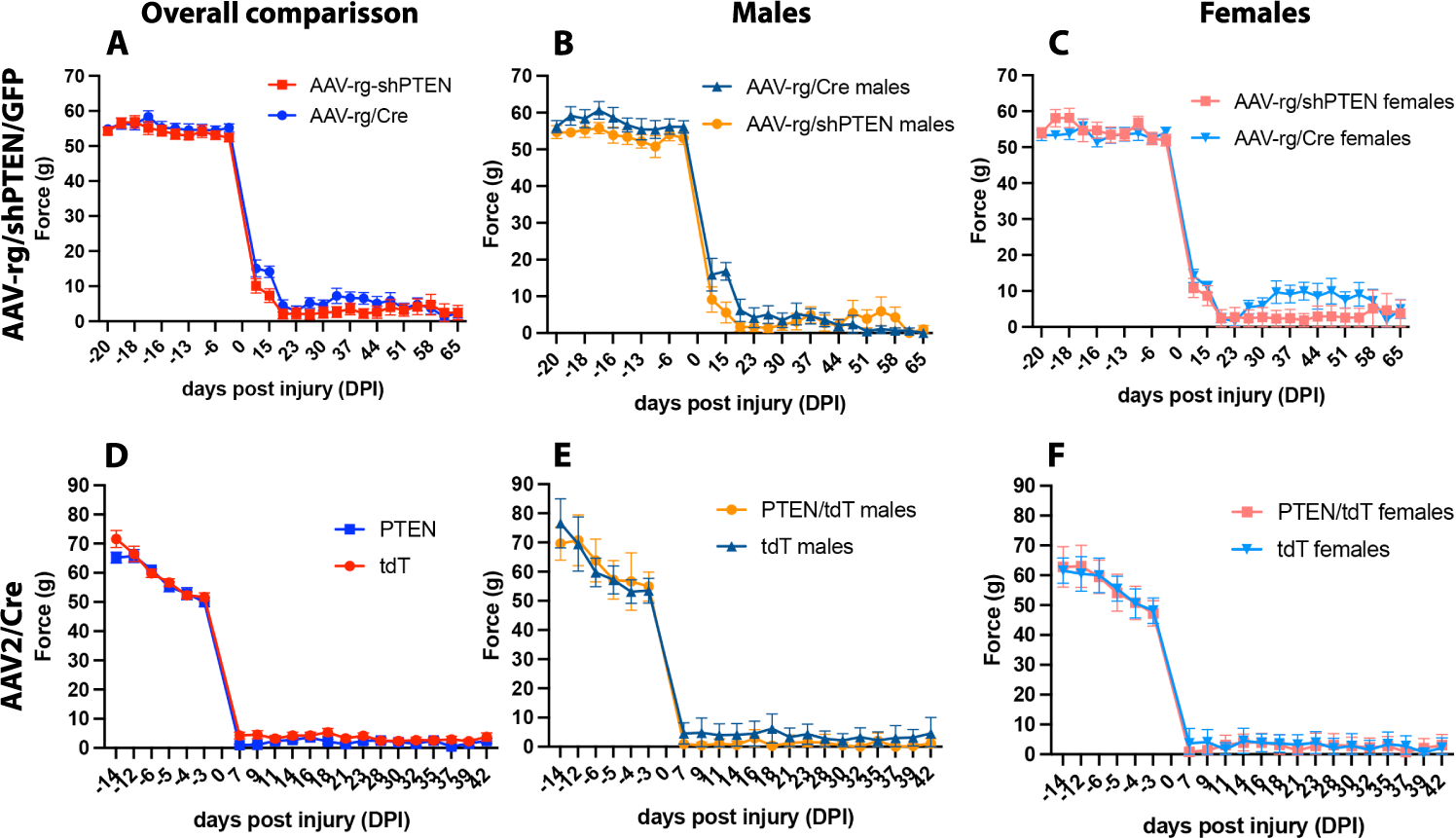
PTEN knockdown with AAV-rg/shPTEN/GFP and PTEN deletion in local spinal circuitry does not promotes functional recovery. Forepaw gripping function as measured by grip strength meter (GSM). A) Rosa^tdTomato^ mice were injected with 3E9 GC/mouse of AAV-rg/shPTEN/GFP or AAV-rg/Cre at the same time as a cervical dorsal hemisection. A mixed effects model analysis revealed significant differences over time [F(535.8) DF 25, 375, p<0.0.001] and but no significant differences between genotypes [F(0.8753) = DF 1, 15, p=0.4301]. B-C) A mixed effects model analysis revealed no significant sex differences for the control Rosa^tdTomato^ mice [F(0.079) DF 1,13, p=0.78] nor for the Rosa^tdTomato^ mice injected with AAV-retro/siPTEN/GFP (PTEN-knockdown) [F(0.11) DF 1,12, p=0.74]. D) Rosa^tdTomato^ and PTEN^f/f^;Rosa^tdTomato^ mice were injected with 6E9 GC/mouse of AAV/Cre. Two-way repeated measures ANOVA revealed significant differences over time [F(988.0) DF 5.13, 123.1, p<0.0001] and no significant differences in grip strength between genotypes [F(4.24) = DF 1,24, p=0.051]. E-F) Two-way repeated measures ANOVA revealed no significant sex differences for the Rosa^tdTomato^ mice [F(3.17) DF 1,10, p=0.11] nor for the PTEN^f/f^;Rosa^tdTomato^ mice [F(0.42) DF 1,12, p=0.53].

##### 3.2.4.a. Comparisons by sex

Stratification of GSM data by sex (Fig. 5B &C) revealed no sex differences in the values of the Rosa^tdTomato^ control mice injected with AAV-rg/Cre and injected with AAV-rg/shPTEN/GFP. GSM values for female mice that received AAV-rg/shPTEN were 4.68% of pre-injury control at the plateau vs. 18.9% for the Rosa^tdTomato^ female control mice. GSM values for male mice that received AAV-rg/shPTEN were 9.57% of pre-injury control at the plateau vs. 4.8% for the Rosa^tdTomato^ male control mice. A mixed effects model analysis revealed no significant sex differences for the control Rosa^tdTomato^ mice [F(0.079) DF 1,13, p=0.78] nor for the Rosa^tdTomato^ mice injected with AAV-rg/shPTEN/GFP (PTEN-knockdown) [F(0.11) DF 1,12, p=0.74]. Overall neither sex showed improvement in recovery of function after SCI plus AAV-rg/shPTEN/GFP injection.

##### 3.2.4.b. Late-developing pathophysiologies

Although mice with AAV-rg/shPTEN/GFP (PTEN knock-down) did not show improvements in motor function in comparison to controls, they did develop pathophysiologies. One male mouse that received AAV-rg/shPTEN/GFP exhibited hindlimb dystonia 1-week post-surgery. This animal was euthanized due to lack of mobility and health concerns. One male mouse that received AAV-rg/shPTEN/GFP exhibited shoulder wounds due to scratching 2.5 weeks post AAV-rg/shPTEN intraspinal injection plus SCI. N=6 mice showed pathophysiologies 5 weeks post AAV-rg/shPTEN/GFP intraspinal injection plus SCI. Of those 6, n=5 (n=3 males; n=2 females) exhibited only dystonia and n=1 (female) exhibited both scratching and dystonia. Of those 6 animals, one mouse was euthanized due to the severity of the scratching wounds. Scratching wounds were treated with Silvadeen and toenails were trimmed. Of n=14 Rosa^tdTomato^ mice (n=6 males; n=8 females), n=8 developed pathophysiologies starting 1 week post AAV-rg/shPTEN/GFP intraspinal injection plus SCI. Of those 8, n=5 were males and n=3 were females; therefore, there was no apparent sex difference in incidence of pathophysiologies with AAV-rg/shPTEN/GFP injections plus SCI. Animals were perfused 12 weeks post AAV-rg/shPTEN/GFP injections plus SCI.

#### 3.2.5. Study 5: PTEN deletion only in local spinal circuitry

Study 5 tested whether pathophysiologies were due to deletion of PTEN in local spinal cord circuitry in the context of a spinal cord injury. Mice received dorsal hemisection injuries and intra-spinal cord injections of AAV2/Cre rather than AAV-rg/Cre (6E9 GC/mouse). Intra-spinal injections of AAV2/Cre delete PTEN in a local region of the spinal cord around the injection site (Fig. 5D) but there is no retrograde transduction of cells of origin of spinal pathways in the brain. PTEN^f/f^;Rosa^tdTomato^ mice received 2 injections above the level of the dorsal hemisection at approximately C4. Controls were Rosa^tdTomato^ mice that received similar injections. Forelimb motor function was assessed over time using a GSM by an individual blinded to treatment group.

Following C5 lesions and AAV2/Cre intra-spinal cord injections, 7 days post-injury GSM values dropped to 8.3% of pre-injury values for Rosa^tdTomato^ mice and 1.8% of pre-injury values for PTEN^f/f^;Rosa^tdTomato^ mice. Values remained low for the duration of the testing period and they plateaued at around 21 days post injury to 5.4% of pre-injury control and to 2.6% of pre-injury values for PTEN^f/f^;Rosa^tdTomato^ mice. Two-way repeated measures ANOVA revealed significant differences over time [F(988.0) DF 5.13, 123.1, p<0.0001] and no significant differences in grip strength between genotypes [F(4.24) = DF 1,24, p=0.051].

##### 3.2.5.a. Comparisons by sex

Stratification of GSM data by sex (Fig. 5E &F) revealed no sex differences in the values of the Rosa^tdTomato^ control mice or in the values of the PTEN^f/f^;Rosa^tdTomato^ mice injected with AAV2/Cre. Specifically, at 21 days post injury, GSM values for female PTEN^f/f^;Rosa^tdTomato^ mice (PTEN-deleted) were 2.7% of pre-injury control at the plateau vs. 7.7% for the Rosa^tdTomato^ female control mice. GSM values for male PTEN^f/f^;Rosa^tdTomato^ mice were 1.7% of pre-injury control at the plateau vs. 1.5% for the Rosa^tdTomato^ male control mice. Two-way repeated measures ANOVA revealed no significant sex differences for the Rosa^tdTomato^ mice [F(3.17) DF 1,10, p=0.11] nor for the PTEN^f/f^;Rosa^tdTomato^ mice [F(0.42) DF 1,12, p=0.53]. Overall neither group showed improvement in recovery of function after SCI plus AAV2/Cre intraspinal injections.

##### 3.2.5.b. Late-developing pathophysiologies

None of the mice that received dorsal hemisections and intra-spinal injections of AAV2/Cre (non-retrograde) exhibited pathophysiologies described above. All animals were euthanized 42 days post injury.

#### 3.2.6. Retrospective analysis of whether there are pathophysiologies with intra-spinal injections of AAV-rg/Cre without concurrent SCI

To assess whether pathophysiologies developed with injections of AAV-rg/Cre without concurrent SCI, we reviewed husbandry and survival records from mice in our previous anatomical studies assessing retrograde transduction via AAV-rg/Cre (Metcalfe et al., 2022; Steward et al., 2021). These studies involved 12 PTEN^f/f^;Rosa^tdTomato^ mice (n=8 males; n=3 females) that were allowed to survive for different intervals after bilateral intra-spinal cord injections of AAV-rg/Cre at C5 (6E9 GC/mouse). Three mice (2 males and 1 female) survived for 1 month post AAV-rg/Cre injection; 4 males survived 4 months post AAV-rg/Cre injection; 1 male survived 7 months post AAV-rg/Cre injection; 5 mice (n=2 males and n=3 females) survived 12 months post AAV-rg/Cre injection. None of these mice exhibited scratching or hindlimb dystonia or any other adverse condition that led to euthanasia.

### 3.3. Transduction of DRG neurons

The incessant scratching that develops suggests pathological itch, which can be due to sensitization of acute itch pathways that originate from defined populations of DRG neurons (Akiyama and Carstens, 2013). Accordingly, we were curious whether DRG neurons that have been implicated in pathological itch are retrogradely transduced. We have previously shown that medium-sized neurons in DRGs within a few segments of the injection site at C5 are retrogradely transduced and that these are not CGRP-positive nociceptors (Metcalfe et al., 2022).

One recent study reported that PTEN deletion in isolectin B4 (IB4)-positive DRG neurons in adult mice, promotes ectopic TRPV1 and Mrgpr3a expression leading to pathological itch (Hu et al., 2022). Sensory representation of the neck and ears is represented at C3, so injections of AAV-rg/Cre at C5 could transduce DRG neurons at C3, deleting PTEN in sensory neurons that innervate the neck and head. This mechanism could account for the incessant scratching we observe if IB4-positive neurons in the DRG are transduced by AAV-rg/Cre injections into the spinal cord.

To rule in or rule out this possible mechanism, we assessed retrograde transduction of DRG neurons from mice in Study 1. Slide-mounted cross-sections from spinal segments near the injection site with DRGs were stained for IB4 and co-imaged for tdT and imaged to assess co-labeling. Many medium sized DRG neurons were tdT positive, there were no convincing examples of co-labeling with IB4 (Fig. 6). This finding does not support the hypothesis that excessive scratching is due to PTEN deletion in IB4-expressing DGR neurons.

**Figure 6.**
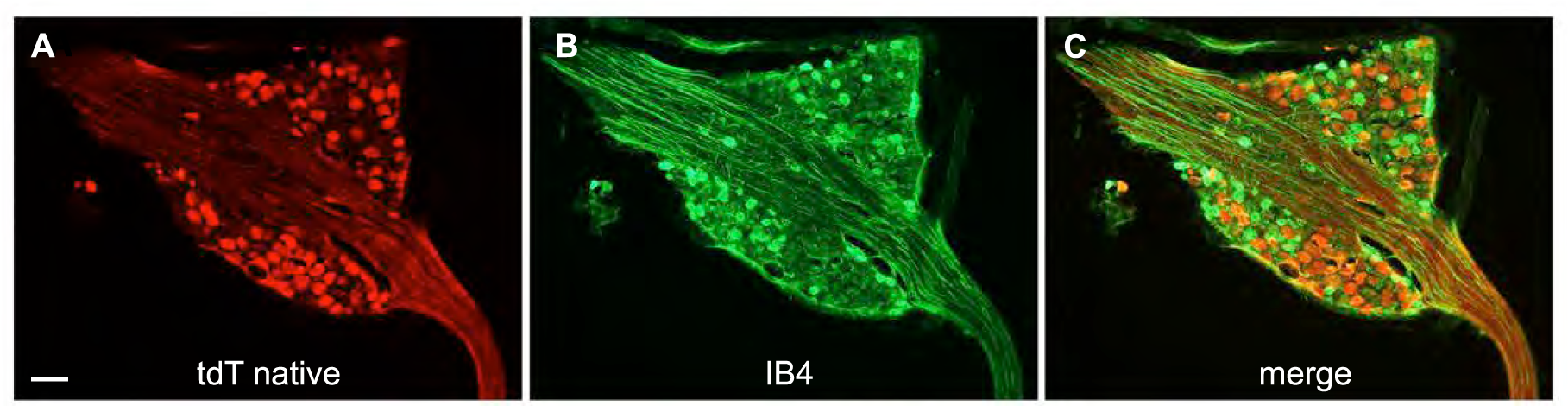
Intraspinal AAV-rg injections transduce DRG neurons with no transduction of IB4+ neurons. A) tdT-positve DRG neurons in PTEN^f/f^;Rosa^tdTomato^ mice. B). IHC for isolectin B4 (IB4)-positive DRG neurons. C) Merge of A) and B) tdT-positive DRG neurons expressing IB4 (if present) would be yellow (none are evident). Scale bar (100 µm).

Another possible circuit for persistent itch involves the mechanical itch pathway (Pan et al., 2019). First order neurons for this circuit are toll-like receptor 5-positive DRG neurons (TLR5+) that give rise to low-threshold mechanoreceptors. These DRG neurons project to a population of Urocortin-3 expressing (Unc3+) excitatory interneurons in the dorsal horn. Thus, a possible mechanism for chronic itch would be increased intrinsic excitability of TLR5+ DRG neurons due to PTEN deletion. As above, this could only occur if this population of DRG neurons is transduced by AAV-rg/Cre injections into the spinal cord. To rule in or rule out this mechanism, slide-mounted cross-sections from DRGs near the injection site with DRGs were stained using what was presumably the same TLR5 antibody from Santa Crus that had been used by (Pan et al., 2019). Although we tried several different immunostaining procedures, we were unable to replicate the findings of Pan et al. in which a select population of DRG neurons were positive for TLR5. Instead, almost all DRG neurons exhibited weak immunofluorescence (data not shown). Because our immunostaining did not reveal a subset of DRG neurons that were selectively labeled for TLR5, it remains possible that some of the DRG neurons transduced by AAV-rg are first order neurons of the mechanical itch pathway.

## 4. DISCUSSION

The overall goal of this study was to evaluate AAV-rg as a potential platform to promote motor recovery after cervical SCI by deleting PTEN in multiple pathways. Our results revealed: 1) Intra-spinal cord injections of AAV-rg/Cre at the time of a dorsal hemisection at C5 in PTEN^f/f^;Rosa^tdTomato^ mice enhanced forelimb motor recovery in a dose (GC) dependent fashion. 2) There were major sex differences in the extent of recovery. Male mice exhibited much greater recovery than females, and the data from males largely accounted for the differences in the combined data. 3) Despite initial recovery, late-onset pathophysiologies involving incessant scratching and hindlimb dystonia developed in many mice, compromising the motor recovery. 4) Development of pathophysiologies was dose-dependent, but pathophysiologies occurred even at low GC doses that did not enhance recovery. 5) Pathophysiologies were also seen with AAV-rg/shPTEN injections at C5, although the GC does used did not enhance motor recovery. 6) Pathophysiologies were not seen with injections of AAV2/Cre at C5 (not retro) or with injections of AAV-rg/Cre without concurrent SCI. In what follows, we discuss implications and caveats of these results.

### 4.1. Dose-dependence for enhanced recovery

Previous studies have reported that PTEN deletion in the sensorimotor cortex can enhance axonal regeneration and functional recovery after spinal cord injury (Danilov and Steward, 2015; Du et al., 2015; Geoffroy et al., 2015; Lewandowski and Steward, 2014; Liu et al., 2010; Zukor et al., 2013). PTEN is a phosphatase that negatively regulates the PI3K/AKT/mTOR signaling pathway, which is crucial for neuronal survival, axonal growth, and regeneration (Leibinger et al., 2019; Proud, 2002; Testa and Tsichlis, 2005; Zhou and Snider, 2006). The present study extends the story by demonstrating that PTEN deletion via retrograde transduction using AAV-rg/Cre can also enhance motor recovery.

Enhanced recovery of forelimb motor function with intra-spinal injections of AAV-rg/Cre at the time of injury was dose-dependent. Recovery was seen at higher doses 12E9 and 6E9 GC (genome copies) and was not seen with the lowest dose tested (3E9 GC). Counts of PTEN-deleted cortical motoneurons (ghost cells in layer V of the sensorimotor cortex) revealed dose-dependent differences in the numbers of PTEN-deleted neurons in Layer V between high and intermediate GC doses.

The likely explanation for the small difference in the numbers of PTEN-negative cells between the high and intermediate GC doses is that the high dose involved 4 injections, 2 rostral and 2 caudal to the injury. Since the dorsal and dorso-lateral CST is completely transected by a dorsal hemisection, injections below the injury would not be expected to transduce many CMNs. One possible explanation for the better outcome in functional recovery at high doses could be that AAV-rg/Cre injections below the injury may be promoting rewiring and synaptic plasticity of intraspinal circuitry. Recent studies have shown that coordinated growth stimulation in supra spinal centers and in spinal relay stations led to enhancement in neuronal rewiring and improved locomotor recovery (Van Steenbergen et al., 2023). The combination of these effects may elicit a greater recovery of function than what is observed with intermediate doses of AAV-rg/Cre.

### 4.3. Sex differences in functional recovery of PTEN-deleted mice after SCI

Our results also demonstrate major sex differences in the response to AAV-rg-mediated PTEN deletion following dorsal hemisection injury in that male mice exhibited much greater recovery than females. Several studies have investigated sex-related differences in functional recovery after SCI in animal models. For instance, female C57Bl/6 mice with thoracic-level compression injuries have shown better locomotor scores and less necrosis and inflammatory cell infiltration than males (Farooque et al., 2006). Additionally, female mice showed significantly better recovery than males after a moderate compression SCI (Hauben et al., 2002). It is worth noting that these studies used compression injuries that can lead to a secondary inflammatory response, which is not as extensive in transection models as used in our study (Watanabe et al., 1999). It is possible that the better outcomes observed in female mice in other studies may be attributed to neuroprotective roles of estrogen-like hormones in the CNS (Yune et al., 2004).

Sex differences in neurologic recovery have been observed in clinical studies of humans, with females showing greater natural recovery than males. However, at the time of discharge from rehabilitation, males tend to perform better functionally for a given level and degree of neurologic injury (Sipski et al., 2004). The underlying reasons for these sex differences are not yet fully understood and could involve a range of factors such as hormone levels, genetics, or other factors. Further studies are needed to elucidate the mechanisms underlying these differences, which could ultimately lead to the development of more effective treatments for spinal cord injury in both males and females.

### 4.4. Late onset pathophysiologies

Despite enhanced motor recovery with AAV-rg-mediated PTEN deletion, many mice exhibited pathophysiologies with all doses used in this study, even low doses that did not elicit functional recovery. Late-onset pathophysiologies occurred only with AAV-rg based delivery of PTEN modifiers (AAV-rg/Cre or AAV-rg/shPTEN/GFP) in conjunction with SCI (dorsal hemisection at C5). This finding suggests several conclusions. 1) Pathophysiologies are not due to PTEN-deletion in spinal circuitry only. This conclusion is supported by the fact that animals that received intraspinal AAV2/Cre injections to delete PTEN in local spinal circuitry (PTEN^f/f^;Rosa^tdT^ + AAV2/Cre + SCI) <u>did not</u> show scratching or hindlimb dystonia. 2) Pathophysiologies are not due to PTEN-deletion alone, since PTEN deleted animals without SCI (PTEN^f/f^;Rosa^tdT^ + AAV-rg/Cre - SCI) do not exhibit pathophysiologies. Although mice that received AAV2/Cre without injury did not exhibit excessive scratching sufficient to cause skin injury, we did not assess scratching behavior directly and so cannot exclude increased scratching with spinal cord PTEN deletion as reported by (Hu et al., 2022).

Some PTEN-deleted animals that received cervical SCI also exhibited rigid forward extension of the hindlimbs (dystonia). Dystonias can be caused by loss of descending inputs. Clinicians use the term “upper motoneuron lesions”, and the explanation is that there is net loss of descending inhibition spinal circuits, resulting in hyperactivity (Bakheit et al., 2011; Sheean and McGuire, 2009). Transduction of CMNs with intraspinal spinal AAV-rg/Cre could disrupt their physiological activity producing functional phenotypes similar to “upper motoneuron lesions”.

Hindlimb dystonias could be also driven by altered connections entirely within the injured spinal cord, again with the constraint that these only occur with PTEN deletion via AAV-rg in conjunction with SCI. The incessant scratching is with the hindlimb, so it is plausible that what begins as an “itch phenotype” converts to hindlimb dystonia driven by the lumbar spinal cord. Future studies will be required to explore these possible mechanisms.

One important clue regarding possible mechanisms is that pathophysiologies are not observed with deletion of PTEN limited to local spinal circuitry (AAV2/Cre rather than AAV-rg/Cre or AAV-rg/shPTEN). This suggests that the pathophysiologies may be due to retrograde transduction and PTEN deletion in neurons that project to the area of the AAV-rg injection. These could include cells of origin any of the descending pathways from the brain, cells of origin of intra-spinal pathways including second order sensory neurons, or first order DRG neurons that give rise to ascending sensory pathways.

A second clue regarding possible mechanisms of the pathophysiologies is that they appear after some delay (4-5 weeks) and increase in incidence and severity over time. This time delay suggests a mechanism that evolves over time, reaching a threshold that triggers aberrant behaviors. Possible explanations include: 1) Growth induced by PTEN deletion leading to the formation of aberrant connections; 2) Persistent activation of the mTOR signaling pathway, which can have profound consequences on the physiological properties of neurons, including causing seizures (Backman et al., 2001; LaSarge et al., 2021; Ogawa et al., 2007; Pun et al., 2012).

### 4.5. Transduction of DRG neurons

In our previous studies involving intra-spinal AAV-rg/Cre injections, we observed retrograde transduction of large DRG neurons in dorsal root ganglia within a few segments of the injection site at C5 (Metcalfe et al., 2022). Sensory representation of the neck and ears is represented at C3, and it is possible that injections of AAV-rg/Cre at C5 transduce some DRG neurons at C3, therefore eliciting PTEN deletion in sensory neurons that innervate the neck and head.

The incessant scratching that develops suggests the possibility of chronic itch in the region of the neck and head. Rodents and other quadrupeds use their hindlimbs to scratch regions of their upper torso, neck, and head to relieve itch or sometimes in response to other sensations. In some pathological conditions, chronic itch is caused by peripheral and central sensitization of acute itch pathways, with sensitization of itch-related somatosensory pathways in DRG neurons (Akiyama and Carstens, 2013).

Although both mechanical and chemical itch trigger site-directed scratching, these two forms are transmitted through different neural circuits. Recent studies have found that interneurons in the dorsal spinal cord expressing urocortin 3 (Ucn3+) (Pan et al., 2019) and neuropeptide Y receptor (Y1R+) (Acton et al., 2019) are essential for transmitting mechanical itch, and they receive inputs from Toll-like receptor 5-positive (TLR5+) Aß low-threshold mechanoreceptors (LTMR). Downstream projection circuits remain unknown. Itch-inducing molecules activate several membrane-bound receptors, such as histamine receptors (Rossbach et al., 2009). Recently, the Mas-related G-protein coupled receptor (Mrgpr) (Liu et al., 2009) and the transient receptor potential (TRP) channel subfamily V member 1 (TRPV1) (Davidson and Giesler, 2010) have been reported to be required for processing the histamine-induced itch response (Davidson and Giesler, 2010). At the spinal level, secondary neurons for pruriceptors express gastrin-releasing peptide (GRP) in the spinal dorsal horn (Bautista et al., 2014; Sun et al., 2017; Sun and Chen, 2007). Information is then transmitted to spinal tertiary pruriceptors expressing the GRP receptor (GRPR). Finally, this information is transmitted to supraspinal regions via a relay of spinal ascending neurons (Mu et al., 2017). It remains to be seen whether any of these neuron types and pathways are transduced by intra-spinal injections of AAV-rg/Cre, and if they are, how consequences could differ in the context of SCI.

Our previous studies with cervical intraspinal injections of AAV-rg/Cre revealed that retrogradely transduced neurons in the DRG (identified by tdT labeling) are distinct from the population neurons that express calcitonin gene-related peptide (CGRP) that mediate nociception (meaning no-co-expression was detected) (Metcalfe et al., 2022). Here, we carried out initial studies to test whether AAV-rg transduced DRG neurons expressing markers of either itch pathway.

To test for transduction of TLR5 neurons, we used what was presumably the same TLR5 antibody from Santa Crus that had been used by Pan et al., 2019. However, we were unable to replicate the findings that a select population of DRG neurons were positive for TLR5. We were unable to determine the RRID for either the antibody used by Pan et al or the one we used, so it is possible that the antibodies from Santa Cruz are from a different batch. Because our immunostaining did not reveal a subset of DRG neurons that were selectively labeled for TLR5, it remains possible that some of the DRG neurons transduced by AAV-rg are first order neurons of the mechanical itch pathway.

Another possible pathway for pathological itch is suggested by the recent finding that PTEN deletion in isolectin B4 (IB4)-positive DRG neurons in adult mice, promotes ectopic TRPV1 and Mrgpr3a expression leading to pathological itch (Hu et al., 2022). This mechanism could account for the incessant scratching we observe if IB4-positive neurons are transduced by AAV-rg/Cre injections into the spinal cord. Our results confirmed the presence of a populaton of IB4-expressing DRG neurons, but there were no convincing examples of retrograde transduction evidenced by co-labeling with tdT. This finding does not support the hypothesis that excessive scratching is due to PTEN deletion in IB4-expressing DGR neurons.

Within the spinal cord, itch signals are transmitted through the spinothalamic tract and spinobrachial pathway to the thalamus and parabrachial nucleus, respectively, where they are projected to various brain areas for itch processing (Yosipovitch et al., 2018). It remains to be seen whether the second order neurons that give rise to these pathways are transduced by AAV-rg. Descending pathways from various sites modulate transmission along sensory pathways, so it is possible that AAV-rg/Cre-mediated PTEN deletion in the cells of origin of these descending pathways alters itch transmission to trigger pathological scratching. Future studies are needed to explore these possible mechanisms.

## 5. CONCLUSION

Our results support the potential of AAV-rg for enhancing motor recovery after SCI, but also highlight the potential for late-developing pathophysiologies with this approach. A recent study reports other indications of late-developing consequences of AAV-rg-mediated PTEN deletion in a thoracic crush model of SCI. Although AAV-rg/Cre-mediated PTEN deletion resulted in significant initial improvement in locomotor function in comparison to control, improvements were not sustained (Stewart et al., 2023). This highlights the importance of tracking potential long-term consequences of manipulations to enhance axon regeneration after SCI. Further studies are necessary to elucidate the mechanisms underlying late-developing adverse effects and functional deterioration and to determine whether recovery can be achieved without adverse consequences.

## Acknowledgements

Supported by NIH grant NS108189 to OS. MM was the recipient of a Career Re-Entry and Diversity Fellowship supplement to NS108189. We gratefully acknowledge generous donations from Cure Medical and Research for Cure. Thanks to Ardi Gunawan for skilled spinal cord surgery, Jamie Mizufuka for superb histological assistance with the immunocytochemistry and analyses, Daniela Gonzalez, Kiara Quinn for assistance in behavioral testing and Kevin Sanchez for assistance in behavioral testing, immunocytochemistry and anatomical analyses.

## Author contributions

OS and MM designed the overall project and wrote the paper.

## REFERENCES

Acton, D., Ren, X., Di Costanzo, S., Dalet, A., Bourane, S., Bertocchi, I., Eva, C., Goulding, M., 2019. Spinal Neuropeptide Y1 Receptor-Expressing Neurons Form an Essential Excitatory Pathway for Mechanical Itch. Cell Rep 28, 625–639.e626.

Akiyama, T., Carstens, E., 2013. Neural processing of itch. Neuroscience 250, 697–714.

Backman, S.A., Stambolic, V., Suzuki, A., Haight, J., Elia, A., Pretorius, J., Tsao, M.S., Shannon, P., Bolon, B., Ivy, G.O., Mak, T.W., 2001. Deletion of Pten in mouse brain causes seizures, ataxia and defects in soma size resembling Lhermitte-Duclos disease. Nat Genet 29, 396–403.

Bakheit, A.M., Fheodoroff, K., Molteni, F., 2011. Spasticity or reversible muscle hypertonia? J Rehabil Med 43, 556–557.

Bautista, D.M., Wilson, S.R., Hoon, M.A., 2014. Why we scratch an itch: the molecules, cells and circuits of itch. Nat Neurosci 17, 175–182.

Challagundla, M., Koch, J.C., Ribas, V.T., Michel, U., Kügler, S., Ostendorf, T., Bradke, F., Müller, H.W., Bähr, M., Lingor, P., 2015. AAV-mediated expression of BAG1 and ROCK2-shRNA promote neuronal survival and axonal sprouting in a rat model of rubrospinal tract injury. J Neurochem 134, 261–275.

Danilov, C.A., Steward, O., 2015. Conditional genetic deletion of PTEN after a spinal cord injury enhances regenerative growth of CST axons and motor function recovery in mice. Exp Neurol 266, 147–160.

Davidson, S., Giesler, G.J., 2010. The multiple pathways for itch and their interactions with pain. Trends Neurosci 33, 550–558.

Du, K., Zheng, S., Zhang, Q., Li, S., Gao, X., Wang, J., Jiang, L., Liu, K., 2015. Pten Deletion Promotes Regrowth of Corticospinal Tract Axons 1 Year after Spinal Cord Injury. J Neurosci 35, 9754–9763.

Farooque, M., Suo, Z., Arnold, P.M., Wulser, M.J., Chou, C.T., Vancura, R.W., Fowler, S., Festoff, B.W., 2006. Gender-related differences in recovery of locomotor function after spinal cord injury in mice. Spinal Cord 44, 182–187.

Gallent, E.A., Steward, O., 2018. Neuronal PTEN deletion in adult cortical neurons triggers progressive growth of cell bodies, dendrites, and axons. Exp Neurol 303, 12–28.

Geoffroy, C.G., Hilton, B.J., Tetzlaff, W., Zheng, B., 2016. Evidence for an Age-Dependent Decline in Axon Regeneration in the Adult Mammalian Central Nervous System. Cell Rep 15, 238–246.

Geoffroy, C.G., Lorenzana, A.O., Kwan, J.P., Lin, K., Ghassemi, O., Ma, A., Xu, N., Creger, D., Liu, K., He, Z., Zheng, B., 2015. Effects of PTEN and Nogo Codeletion on Corticospinal Axon Sprouting and Regeneration in Mice. J Neurosci 35, 6413–6428.

Guertin, D.A., Sabatini, D.M., 2007. Defining the role of mTOR in cancer. Cancer Cell 12, 9–22.

Hauben, E., Mizrahi, T., Agranov, E., Schwartz, M., 2002. Sexual dimorphism in the spontaneous recovery from spinal cord injury: a gender gap in beneficial autoimmunity? Eur J Neurosci 16, 1731–1740.

Hu, L., Jiang, G.Y., Wang, Y.P., Hu, Z.B., Zhou, B.Y., Zhang, L., Song, N.N., Huang, Y., Chai, G.D., Chen, J.Y., Lang, B., Xu, L., Liu, J.L., Li, Y., Wang, Q.X., Ding, Y.Q., 2022. The role of PTEN in primary sensory neurons in processing itch and thermal information in mice. Cell Rep 39, 110724.

Kaplitt, M.G., Feigin, A., Tang, C., Fitzsimons, H.L., Mattis, P., Lawlor, P.A., Bland, R.J., Young, D., Strybing, K., Eidelberg, D., During, M.J., 2007. Safety and tolerability of gene therapy with an adeno-associated virus (AAV) borne GAD gene for Parkinson’s disease: an open label, phase I trial. Lancet 369, 2097–2105.

LaSarge, C.L., Pun, R.Y.K., Gu, Z., Riccetti, M.R., Namboodiri, D.V., Tiwari, D., Gross, C., Danzer, S.C., 2021. mTOR-driven neural circuit changes initiate an epileptogenic cascade. Prog Neurobiol 200, 101974.

Leibinger, M., Hilla, A.M., Andreadaki, A., Fischer, D., 2019. GSK3-CRMP2 signaling mediates axonal regeneration induced by Pten knockout. Commun Biol 2, 318.

Lewandowski, G., Steward, O., 2014. AAVshRNA-mediated suppression of PTEN in adult rats in combination with salmon fibrin administration enables regenerative growth of corticospinal axons and enhances recovery of voluntary motor function after cervical spinal cord injury. J Neurosci 34, 9951–9962.

Liu, K., Lu, Y., Lee, J.K., Samara, R., Willenberg, R., Sears-Kraxberger, I., Tedeschi, A., Park, K.K., Jin, D., Cai, B., Xu, B., Connolly, L., Steward, O., Zheng, B., He, Z., 2010. PTEN deletion enhances the regenerative ability of adult corticospinal neurons. Nat Neurosci 13, 1075–1081.

Liu, Q., Tang, Z., Surdenikova, L., Kim, S., Patel, K.N., Kim, A., Ru, F., Guan, Y., Weng, H.J., Geng, Y., Undem, B.J., Kollarik, M., Chen, Z.F., Anderson, D.J., Dong, X., 2009. Sensory neuron-specific GPCR Mrgprs are itch receptors mediating chloroquine-induced pruritus. Cell 139, 1353–1365.

Metcalfe, M., Yee, K.M., Luo, J., Martin-Thompson, J.H., Gandhi, S.P., Steward, O., 2022. Harnessing rAAV-retro for gene manipulations in multiple pathways that are interrupted after spinal cord injury. Exp Neurol 350, 113965.

Mu, D., Deng, J., Liu, K.F., Wu, Z.Y., Shi, Y.F., Guo, W.M., Mao, Q.Q., Liu, X.J., Li, H., Sun, Y.G., 2017. A central neural circuit for itch sensation. Science 357, 695–699.

Ogawa, S., Kwon, C.H., Zhou, J., Koovakkattu, D., Parada, L.F., Sinton, C.M., 2007. A seizure-prone phenotype is associated with altered free-running rhythm in Pten mutant mice. Brain Res 1168, 112–123.

Pan, H., Fatima, M., Li, A., Lee, H., Cai, W., Horwitz, L., Hor, C.C., Zaher, N., Cin, M., Slade, H., Huang, T., Xu, X.Z.S., Duan, B., 2019. Identification of a Spinal Circuit for Mechanical and Persistent Spontaneous Itch. Neuron 103, 1135–1149.e1136.

Park, K.K., Liu, K., Hu, Y., Kanter, J.L., He, Z., 2010. PTEN/mTOR and axon regeneration. Exp Neurol 223, 45–50.

Park, K.K., Liu, K., Hu, Y., Smith, P.D., Wang, C., Cai, B., Xu, B., Connolly, L., Kramvis, I., Sahin, M., He, Z., 2008. Promoting axon regeneration in the adult CNS by modulation of the PTEN/mTOR pathway. Science 322, 963–966.

Proud, C.G., 2002. Regulation of mammalian translation factors by nutrients. Eur J Biochem 269, 5338–5349.

Pun, R.Y., Rolle, I.J., Lasarge, C.L., Hosford, B.E., Rosen, J.M., Uhl, J.D., Schmeltzer, S.N., Faulkner, C., Bronson, S.L., Murphy, B.L., Richards, D.A., Holland, K.D., Danzer, S.C., 2012. Excessive activation of mTOR in postnatally generated granule cells is sufficient to cause epilepsy. Neuron 75, 1022–1034.

Rossbach, K., Wendorff, S., Sander, K., Stark, H., Gutzmer, R., Werfel, T., Kietzmann, M., Bäumer, W., 2009. Histamine H4 receptor antagonism reduces hapten-induced scratching behaviour but not inflammation. Exp Dermatol 18, 57–63.

Schneider, C.A., Rasband, W.S., Eliceiri, K.W., 2012. NIH Image to ImageJ: 25 years of image analysis. Nat Methods 9, 671–675.

Sheean, G., McGuire, J.R., 2009. Spastic hypertonia and movement disorders: pathophysiology, clinical presentation, and quantification. Pm r 1, 827–833.

Sipski, M.L., Jackson, A.B., Gómez-Marín, O., Estores, I., Stein, A., 2004. Effects of gender on neurologic and functional recovery after spinal cord injury. Arch Phys Med Rehabil 85, 1826–1836.

Steward, O., Yee, K.M., Metcalfe, M., Willenberg, R., Luo, J., Azevedo, R., Martin-Thompson, J.H., Gandhi, S.P., 2021. Rostro-Caudal Specificity of Corticospinal Tract Projections in Mice. Cereb Cortex 31, 2322–2344.

Stewart, A.N., Kumari, R., Bailey, W.M., Glaser, E.P., Hammers, G.V., Wireman, O.H., Gensel, J.C., 2023. PTEN knockout using retrogradely transported AAVs restores locomotor abilities in both acute and chronic spinal cord injury. bioRxiv, 2023.2004.2017.537179.

Sun, S., Xu, Q., Guo, C., Guan, Y., Liu, Q., Dong, X., 2017. Leaky Gate Model: Intensity-Dependent Coding of Pain and Itch in the Spinal Cord. Neuron 93, 840–853.e845.

Sun, Y.G., Chen, Z.F., 2007. A gastrin-releasing peptide receptor mediates the itch sensation in the spinal cord. Nature 448, 700–703.

Tervo, D.G., Hwang, B.Y., Viswanathan, S., Gaj, T., Lavzin, M., Ritola, K.D., Lindo, S., Michael, S., Kuleshova, E., Ojala, D., Huang, C.C., Gerfen, C.R., Schiller, J., Dudman, J.T., Hantman, A.W., Looger, L.L., Schaffer, D.V., Karpova, A.Y., 2016. A Designer AAV Variant Permits Efficient Retrograde Access to Projection Neurons. Neuron 92, 372–382.

Testa, J.R., Tsichlis, P.N., 2005. AKT signaling in normal and malignant cells. Oncogene 24, 7391–7393.

Van Steenbergen, V., Burattini, L., Trumpp, M., Fourneau, J., Aljović, A., Chahin, M., Oh, H., D’Ambra, M., Bareyre, F.M., 2023. Coordinated neurostimulation promotes circuit rewiring and unlocks recovery after spinal cord injury. J Exp Med 220.

Wang, Z., Maunze, B., Wang, Y., Tsoulfas, P., Blackmore, M.G., 2018. Global Connectivity and Function of Descending Spinal Input Revealed by 3D Microscopy and Retrograde Transduction. J Neurosci 38, 10566–10581.

Watanabe, T., Yamamoto, T., Abe, Y., Saito, N., Kumagai, T., Kayama, H., 1999. Differential activation of microglia after experimental spinal cord injury. J Neurotrauma 16, 255–265.

Willenberg, R., Zukor, K., Liu, K., He, Z., Steward, O., 2016. Variable laterality of corticospinal tract axons that regenerate after spinal cord injury as a result of PTEN deletion or knock-down. n/a-n/a.

Yosipovitch, G., Rosen, J.D., Hashimoto, T., 2018. Itch: From mechanism to (novel) therapeutic approaches. J Allergy Clin Immunol 142, 1375–1390.

Yune, T.Y., Kim, S.J., Lee, S.M., Lee, Y.K., Oh, Y.J., Kim, Y.C., Markelonis, G.J., Oh, T.H., 2004. Systemic administration of 17beta-estradiol reduces apoptotic cell death and improves functional recovery following traumatic spinal cord injury in rats. J Neurotrauma 21, 293–306.

Zhou, F.Q., Snider, W.D., 2006. Intracellular control of developmental and regenerative axon growth. Philos Trans R Soc Lond B Biol Sci 361, 1575–1592.

Zukor, K., Belin, S., Wang, C., Keelan, N., Wang, X., He, Z., 2013. Short hairpin RNA against PTEN enhances regenerative growth of corticospinal tract axons after spinal cord injury. J Neurosci 33, 15350–15361.

